# Graph-based characterization of in vitro neuronal network maturation using machine learning and digital holographic microscopy

**DOI:** 10.64898/2026.06.18.732973

**Authors:** Zahra Yazdani, Erik Bélanger, Maxime Moreaud, Jodie Llinares, Antoine Allard, Pierre Marquet, Patrick Desrosiers

## Abstract

**Significance:** Digital Holographic Microscopy (DHM) provides label-free quantitative phase images (QPIs) of living cells and has become a powerful tool for studying cellular morphology and dynamics. While most DHM studies have focused on cell-level analysis, the quantitative characterization of neuronal network organization and maturation from DHM images remains largely unexplored, highlighting the need for dedicated computational approaches.

**Aim:** We aimed to develop an automated framework combining deep-learning-based image analysis and graph theory to quantitatively characterize the organization, connectivity, and maturation of neuronal networks in primary rat cortical cultures imaged by DHM.

**Approach:** Two U-Net convolutional neural networks were trained on manually annotated DHM phase images to segment neuronal cell bodies and neurites. The resulting segmentation maps were used to infer putative morphological connections between neurons and generate graph representations of neuronal networks, referred to as graph fingerprints. A panel of 18 connectomics-inspired graph features was then computed to characterize local and global properties of network organization across four stages of culture maturation.

**Results:** The mean area under the receiver operating characteristic curves was 0.98 for cell-body and 0.91 for neurite segmentation, indicating near-perfect identification. Graph-theoretical analysis revealed reproducible topological changes during network maturation in vitro, including increased density, reduced modularity, and progressive network integration. Correlation analysis showed that the 18 graph features grouped into two highly correlated families. A Random Forest classifier identified density and modularity as the most informative descriptors, achieving an accuracy of 87% in classifying maturation stages of neuronal cultures.

**Conclusions:** Our results demonstrate that combining DHM, deep-learning-based segmentation, and graph-theoretical analysis enables quantitative characterization of neuronal network organization and maturation from label-free phase images. This framework provides a foundation for future studies of pharmacological experiments, neuronal network phenotyping, and human induced pluripotent stem cell (hiPSC)-derived neuronal cultures, where quantitative assessment of network organization remains a major challenge.

## 1 Introduction

Over the past two decades, quantitative phase imaging (QPI) has advanced significantly, becoming a widely used optical microscopy technique for label-free live-cell imaging.^1^ Biological cells behave predominantly as weakly scattering phase objects, making phase-sensitive imaging particularly attractive since phase provides an intrinsic contrast mechanism for transparent specimens. While classical phase-contrast and differential interference contrast (DIC) microscopy provide high-quality visualization of transparent specimens, they remain largely qualitative and may introduce imaging artifacts that complicate image interpretation and automated image analysis.

Among QPI techniques, Digital Holographic Microscopy (DHM) occupies a unique position by combining interferometric sensitivity with computational wavefront reconstruction. In off-axis configurations, both the amplitude and phase of the optical field can be retrieved from a single hologram without mechanical scanning.^2^ Because holograms are recorded out of focus and numerically propagated during reconstruction, the diffracted wavefront can be refocused to arbitrary planes, enabling correction of optical aberrations and experimental drifts as well as extended depth of field.^3^ These computational imaging capabilities, together with the high phase sensitivity provided by interferometric detection, make DHM particularly well suited for long-term label-free imaging of living cells and their morphology, including cell bodies and fine cellular processes.^4^

A key output of DHM is the quantitative phase signal (QPS), which reflects the optical path delay (OPD) induced by the specimen and depends on both cell thickness and refractive index contrast.^5^ Most DHM studies have exploited the QPS to derive OPD-based morphological and mass-related descriptors at the level of individual cells, including cell morphology, dry mass, growth dynamics, cell-cycle progression, and viability assessment.^6–10^ Beyond these biophysical measurements, recent advances in machine learning and artificial intelligence have enabled the extraction of complex OPD-derived phenotypic signatures for cell classification, state identification, and diagnostic applications.^11–14^ Combined with high-content phenotyping, these approaches have also facilitated the monitoring of cellular responses to perturbations such as drug exposure, further expanding the scope of label-free quantitative cell analysis.^15–17^ Despite these advances, the potential of DHM for studying neuronal networks remains largely underexploited. By enabling label-free visualization of neuronal architecture over large fields of view, DHM offers a unique opportunity to investigate the structural organization and maturation of neuronal networks.^4^

Although various analysis tools exist for microscopy images, none adequately addresses the demands of large-scale analysis and the unique properties of DHM quantitative phase images. Existing approaches range from manual and semi-automatic annotation tools to fully automated deeplearning methods, each presenting important limitations when applied to DHM neuronal images. Non-automatic methods, such as thresholding, and semi-automatic toolboxes in Fiji’s plugins,^18^ like Labkit,^19^ Simple Neurite Tracer (SNT),^20^ and Trainable Weka Segmentation,^21^ offer quantification of neuronal anatomy but are impractical for large microscopy datasets. Furthermore, fully automatic methods, such as DeepNeurite^22^ for neurite segmentation and Cellpose^23^ for cell body annotation, face challenges in separating cytoplasm from nuclei and distinguishing neurite branches near cell bodies in DHM phase images because the QPS is both non-specific and axially cumulative.^24^

To address the need for enhanced analysis tools, we introduce a novel computational framework designed for the automated and precise analysis of DHM phase images of primary rat cortical neuronal cultures. In line with the increasing adoption of machine learning for microscopy image segmentation tasks,^25^ our framework employs two U-Net convolutional neural network (CNN) models:^26–28^ one dedicated to cell body segmentation and the other to neurite segmentation. These models are trained on manually annotated DHM phase images of neurons, enabling accurate delineation of cell bodies and neurites for detailed analysis of neuronal network structure. After completing the segmentation, we apply a pathfinding algorithm to construct a graph model for each phase image, referred to as the graph fingerprint of primary rat cortical neuronal cultures. To facilitate a comprehensive investigation of network properties, we analyze a set of 18 connectomics-inspired graph features.^29–34^ These features provide insights into both local and global characteristics of primary rat cortical neuronal cultures, enabling the identification of distinct neurodifferentiation stages and structural properties from a network neuroscience perspective.^35–37^

## 2 Materials and methods

### 2.1 Neuronal cell culture

We obtain primary cortical neurons from the brain cortex of neonatal rats (P_0_-P_1_), as previously described.^4^ After dissection, we dissociate the cells enzymatically using 12 U/mL papain (LS003127, Worthington Biochemical) and mechanically by trituration with a Pasteur pipette. We centrifuge and then plate the cells at an average density of 135000 cells on 18-mm diameter round borosilicate coverslips coated with poly-D-lysine (GG18-1.5-PDL, Neuvitro). We maintain the samples in neurobasal media supplemented with B27 (LS21103049, Gibco), 50 U/mL penicillin (15140122, Life Technologies), 50 µg/mL streptomycin (15140122, Life Technologies), 0.5 mM L-glutamax (35050-061, Life Technologies), 5% heat-inactivated fetal bovine serum (SH3007902, Fisher Scientific), and 5 µM cytosine arabinofuranoside (C1768-100MG, Sigma) between day 5 and day 9. Then, we culture and plate neurons at 37°C in a humidified atmosphere of 95% air and 5% CO_2_ for up to 21 days in vitro (DIV), with medium changes twice a week. The primary rat cortical neuronal cell cultures preparation steps are shown in Fig. 1-A.

**Fig 1.**
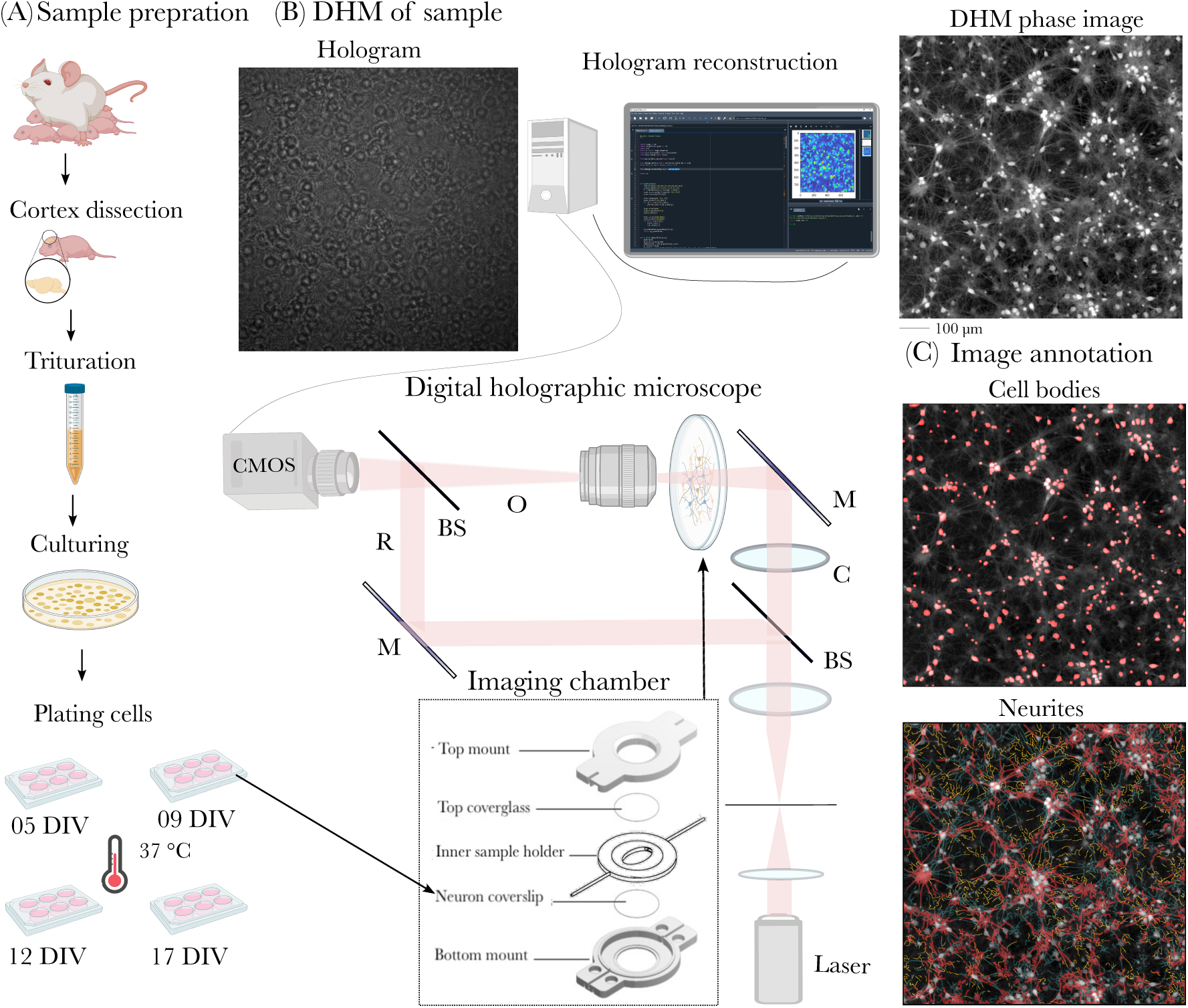
Imaging pipeline. **(A)** Sample preparation: Rat cortices from postnatal (P0-P1) rats are dissected, triturated, cultured, and plated on coverslips in a 37°C incubator. **(B)** DHM of samples: Coverslips are mounted in the DHM imaging chamber. A laser beam passes through a beam splitter (BS), a condenser (C), and a mirror (M), to the sample. Object (O) and reference (R) waves interfere on the CMOS sensor, forming a hologram and DHM phase image is reconstructed. (C) Image annotation: Cell bodies (top) and neurites (bottom) are manually annotated.

### 2.2 Digital holographic microscopy

We use a Mach-Zehnder-based digital holographic microscope (T-1003, Lyncée Tec) with a 666-nm laser diode. The emitted beam is split into two, forming an object wave and a reference wave. The object wave is scattered by the sample and collected by a microscope objective interferes with the reference wave, generating an off-axis hologram. We employ a microscope objective with a magnification of 5x (HC PL FLUOTAR, Leica) which have a working distance of 13.7 mm and a numerical aperture of 0.15. The off-axis geometry results in a small angle between the propagation directions of the two waves. The holograms with a resolution of 1024×1024 pixels are captured using a digital camera equipped with a 2.3-megapixel monochrome CMOS sensor with a pixel size of 5.86 µm and a frame rate of 164 fps (acA1920-155um, Basler). We integrate digital holograms over an interval of 0.5 ms and acquire them at a frequency of 10 Hz for a duration of 10 s. We combine these 100 phase images using maximum intensity projection to produce a single, slightly denoised, detailed phase image that captures comprehensive structural features of the neuronal network of primary rat cortical cell cultures.^38^ The microscope setup is shown in Fig. 1-B.

We reconstruct phase images from holograms by simulating the illumination of the recorded hologram with a digital reference wave using a custom-made reconstruction pipeline consisting of five steps.^39^ First, we filter the hologram in the frequency domain to isolate one of the imaging terms (also known as +1 order of diffraction, or virtual image). In the second step, we compensate for various phase aberrations, such as the tilt introduced by off-axis geometry and spherical aberration from the microscope objective, by positioning a numerical parametric lens (also known as a digital phase mask) in the hologram plane. At the third step, we automatically determine the reconstruction distance corresponding to the best plane of focus. Next, we propagate the corrected object wave to the said reconstruction plane using the convolution approach. Finally, we extract the phase from the reconstructed object wave to form a phase image with a resolution of 800×800 pixels, resulting in a field of view (FOV) of 937.6 µm × 937.6 µm.

### 2.3 Longitudinal live-cell imaging protocol

We keep neurons humidified at 37°C in an incubator until imaging, after which they are transferred to a 3D-printed imaging chamber.^40^ We then fill the chamber with artificial cerebrospinal fluid, which has the following composition (in mM) 100 NaCl, 5.4 KCl, 1.8 CaCl_2_, 0.8 MgCl_2_, 0.9 NaH_2_PO_4_, 10 HEPES, 5 D_glucose_ (pH 7.4; 225-230 mOsm). The imaging chamber is shown in Fig. 1-B.

We performed imaging on primary rat cortical neuronal cell cultures at four time points: 5, 9, 12, and 17 DIV. These time points were selected to capture major transitions in structural maturation and connectivity organization. Previous electrophysiological studies have shown that neuronal cultures exhibit progressively increasing firing and bursting activity as they mature over time.^41^ In parallel, during the first week in vitro, neuronal cultures undergo extensive neuritic outgrowth and branching, leading to increased network complexity and intercellular connectivity.^42–45^ Around 10 DIV, cortical neurons form increasingly elaborate and interconnected networks, whereas at later stages, structural alterations and neuritic degradation may progressively emerge.^43^ Together, these selected time points enable the assessment of key structural changes in neuronal cultures over time. We image a total of nearly 100 coverslips from three different batches of sample preparation. For each of these coverslips, we acquire up to four different FOVs. From this extensive data collection, we specifically select 192 DHM phase images and each of the maturation stages is represented by 48 phase images. This careful selection and stratification ensured a balanced representation across the maturation timeline, a crucial point for analyzing the structural evolution and network connectivity of neuronal networks. A typical phase image is shown in Fig. 1-B top-left.

### 2.4 Labeling phase images

We select a total of 32 phase images from the different maturity stages and manually segment cell bodies using the Cellpose GUI.^23^ Through this process, we obtain accurate binary mask images of cell bodies (red) for all 32 phase images, one of which is shown in Fig. 1-C top. In addition, we use Labkit^19^ to manually annotate neuronal connections on 6 phase images. For ease of annotation, neuronal processes are categorized into three classes: narrow, medium, and wide connections. As shown in Fig. 1-C bottom, each category is labeled using three different brush sizes: 1 pixel (gold), 2 pixels (dark green), and 4 pixels (brown), specifically curated to segment axonal projections, dendrites, and other patterns essential for maintaining neuronal connectivity. In total, the process of annotating all the cell bodies and neuronal processes take approximately 220 hours of work. More details can be found in the Supplementary Material in Section A.

### 2.5 U-Net architecture and training

We use two U-Net models with the same architecture but trained separately: one for cell bodies and one for neurites. Each implemented U-Net model is based on the work of Ronneberger et al.,^26, 27^ adapted for Keras.^28^ The architecture of the U-Net models consists of an encoder-decoder structure, as shown in the middle of Fig. 2. Numbers below each layer represent the number of features, while those to the right indicate pixel dimensions. The final layer of the models is a sigmoid function that provides a probability map image. More details can be found in the Supplementary Material in Section B.

**Fig 2.**
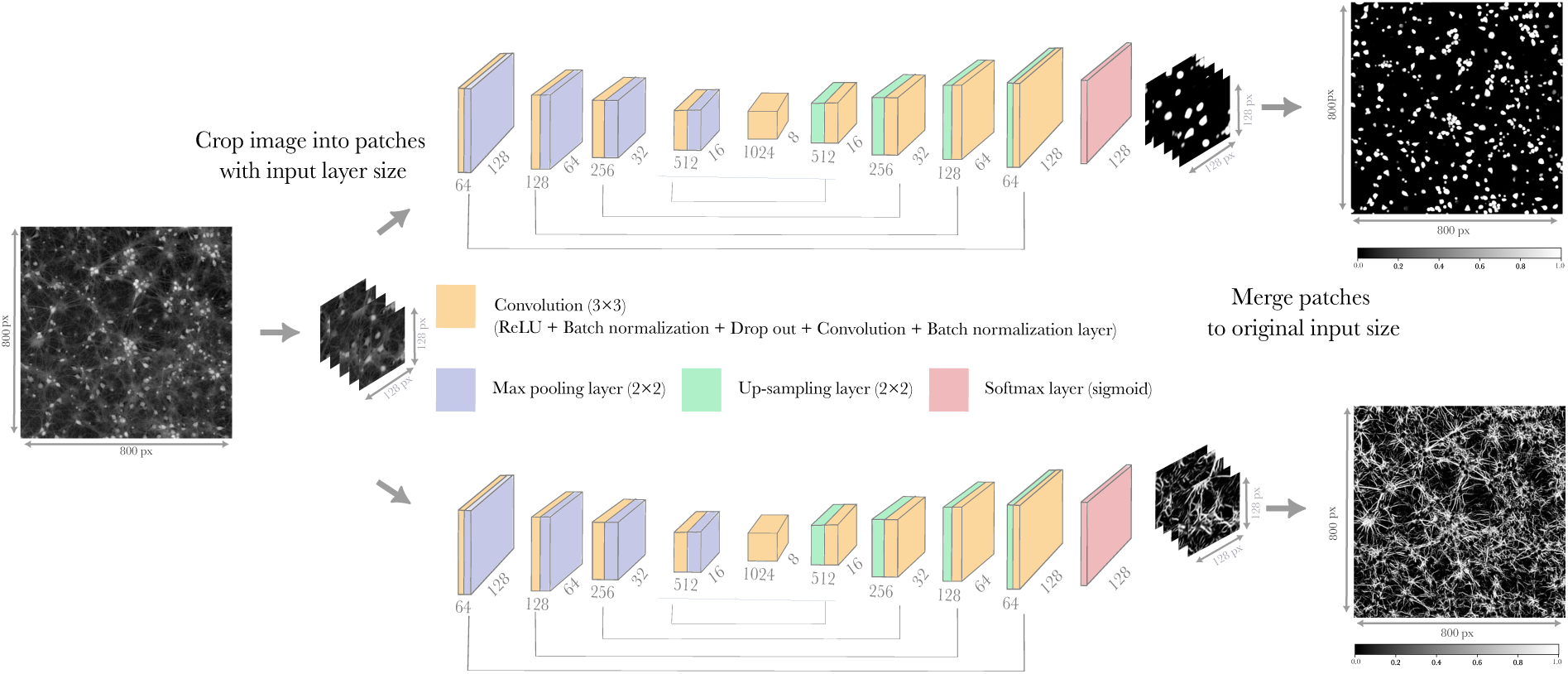
Segmentation pipeline using two U-Net models with a patch-wise approach. The input, shown on the left, is a grayscale image with values between 0 and 1. It is first divided into smaller patches, which serve as inputs to the U-Net models. The U-Net pipeline is shown in the middle, with numbers below each layer representing features and those to the right indicating pixel dimensions. Using a stochastic patch-wise method, the output probability map patches are merged to generate two complete probability maps—one for cell bodies and one for neurites—matching the original image size. The right side of the figure displays the final probability maps, where pixel values range from 0 to 1, representing the predicted likelihood of each pixel belonging to a cell body (trop) or a neurite (bottom).

For training, we select 30 out of 32 images with manually segmented cell bodies and 5 out of 6 images with manually segmented neurites, with the remaining images used for testing model performance. We randomly crop the full-size images into 128×128-pixel patches, matching the input layer size of the U-Net models. To enhance the training dataset, we apply real-time data augmentation to the input images, improving model robustness and generalization by simulating various transformations. These transformations include rotations, random shifts in horizontal and vertical directions, shearing, zooming, and horizontal mirroring. Any missing pixels resulting from these transformations are filled using the nearest-neighbor algorithm. This augmentation process expands the training dataset 20-fold, helping to prevent overfitting and enabling the model to learn more invariant and generalized features. We use 80% of the patches for training and 20% for validation. Each model is trained for 50 epochs, and training is stopped when performance converged between the training and validation sets.

In order to use the trained models, the segmentation pipeline is complemented by patch-extracting and patch-recombining modules, as illustrated on the left and right of Fig. 2. To adapt the original full-size images to the U-Net input layer, we use a stochastic patch-wise method.^46^ This method divides each full-size image into patches while randomly assigning positions to them, helping to reduce edge effects compared to traditional methods like patchify^47^ or empatches,^48^ which simply split images into overlapping patches. Each patch is processed individually by the U-Net models, and the outputs are then merged using the same stochastic patch-wise method to reconstruct full-size probability maps.

### 2.6 U-Net models performance assessment

We evaluate the performance of our two U-Net models by comparing their predicted probability maps with ground truth binary masks of test images. Each probability map is thresholded to enable direct comparison with the binary ground truth masks: pixels above the threshold are classified as objects, while those below are classified as background. Based on this classification, each pixel falls into one of the following categories:

1. True positives (TP): The model correctly predicts an object pixel as an object;
2. False negatives (FN): The model fails to identify an object pixel as an object;
3. True negatives (TN): The model correctly predicts a background pixel as a background;
4. False positives (FP): The model incorrectly identifies a background pixel as an object.

Using these categories, we compute standard segmentation performance metrics. The true positive rate (TPR, or sensitivity) measures the proportion of actual object pixels (cell bodies or neurites) correctly identified by the model, 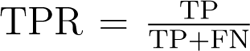, while the false positive rate (FPR) quantifies the proportion of background pixels incorrectly classified as objects, 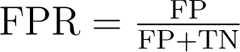

A key part of the performance evaluation involves constructing the receiver operating characteristic (ROC) curve. For this, we use two full-size test images of cell bodies and one full-size test image of neurites. Each full-size test image is cropped into 100 patches, which are individually segmented by the corresponding U-Net model to generate probability maps. To build the ROC curve, we systematically vary the threshold applied to the probability maps, sweeping from 0 to 1. At each threshold, the TPR and FPR are calculated. This process generates a set of (TPR, FPR) pairs, which form the ROC curve. Finally, we assess model performance by computing the area under the curve (AUC) from the average ROC curve, obtained by averaging the individual ROC curves of the test images for each segmentation task.

### 2.7 Optimal threshold selection and final image binarization

We determine the optimal threshold from the ROC curve analysis by identifying the value that maximizes either the F-score or Balanced Accuracy. The F-score and Balanced Accuracy are calculated as 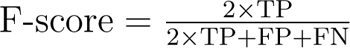 and 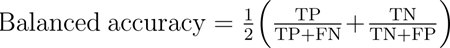 respectively. The optimal threshold is then applied to the probability maps of the U-Net models to generate binary masks. Each pixel is classified as either background or foreground, corresponding to cell bodies or neurites. To separate possible touching cell bodies in these binary masks, we use a watershed algorithm from the scikit-image package.^49^ Finally, for each input phase image we obtain binary masks for cell bodies and neurites, with foreground areas coded as 1 and background as 0.

### 2.8 Graph inference

In graph theory, a graph is defined by its elements (nodes) and their pairwise connections (edges), often represented in an adjacency matrix.^50^ In this article, each node represents a cell body, and each edge represents a neurite likely connecting two cell bodies. This structural information forms the basis of what we call a graph fingerprint. The graph fingerprint serves as an abstract representation of the underlying neuronal network imaged by DHM, encapsulating its most salient features.

To infer the graph fingerprint of a neuronal network, we use the binary mask images of the corresponding cell bodies and neurites, along with a pathfinding algorithm inspired by bidirectional search algorithms.^51^ The first step of the process consists in combining the two mask images with a Boolean OR operator to create a single segmentation mask. This segmentation mask is then transformed into a maze-like structure where values represented different elements: −1 for walls, 0 for passable points containing neurites and cell bodies, and an integer *k >* 0 for centroids of the *k*-th cell body, which serves as node *k* in the graph. To determine possible connections between nodes, an iterative propagation process is used. Starting from each node *k* (cell body centroid), the corresponding *k*-value spreads outward to adjacent 0-valued pixels, incrementing the recorded distance from the source by one at each iteration. This progression continues until different paths meet, meaning *k*-values from different sources overlap. The first intersection between two paths is recorded, as it corresponds to the shortest path between two nodes along passable regions, thereby forming an edge in the graph, representing a potential connection between two cell bodies.

### 2.9 Graph-theoretic features

The procedure presented in the previous section returns, for each imaged neuronal network, a graph fingerprint in the form of an undirected, unweighted graph. To quantitatively compare these graphs, particularly across maturation stages, we systematically characterize their topology, which encompasses their structural organization and connectivity patterns. For this, we select a comprehensive array of 18 graph-theoretic features inspired by connectome analysis at different spatial scales,^29, 31, 33, 35^ which are listed in Table 1.

**Table 1.** List of graph features used for the analysis.

|  |  |
| --- | --- |
| 1. # of connected components | 10. FONs in the LCC center |
| 2. FONs in the LCC | 11. FONs in the LCC periphery |
| 3. Connected components entropy | 12. FONs in the LCC barycenter |
| 4. Density | 13. LCC efficiency |
| 5. LCC mean assortativity coefficient | 14. Average clustering coefficient |
| 6. LCC average distance | 15. Average squares clustering coefficient |
| 7. Average node betweenness centrality | 16. Transitivity |
| 8. Average edge betweenness centrality | 17. # of communities |
| 9. LCC average eccentricity | 18. Modularity |

We first compute the *number of connected components*, where each component is a maximal subset of nodes in which any two nodes are connected by a sequence of edges. These components range from isolated nodes to small and large clusters of connected nodes. Among all components, we then identify the *largest connected component (LCC)*, which corresponds to the largest set of nodes that are all linked, either directly or indirectly, through connections. This allows us to measure the *fraction of nodes (FONs) in the LCC*. To assess how evenly nodes are distributed across components, we compute the *connected components entropy* using the normalized Shannon entropy. A value close to 0 indicates that most nodes belong to a single component, while values near 1 suggest a more even distribution across multiple components.

Several of the selected features are related to node degree, which quantifies the number of connections a node has. These include graph *density*, which represents the proportion of existing connections relative to the total possible connections, and the *LCC mean assortativity coefficient*, which assesses the tendency of nodes in the LCC to connect with others of similar degree.

Other features involve paths within the graph, i.e., sequences of edges connecting two nodes. A shortest path is the minimal sequence of edges linking two distinct nodes, with its length defined as the number of edges in the sequence. The *LCC average distance* quantifies the mean shortest path length between all pairs of nodes within the LCC. A node’s betweenness centrality measures the fraction of shortest paths that pass through it. Nodes with high betweenness centrality values therefore participate in a large number of shortest paths. Similarly, an edge’s betweenness centrality measures the fraction of shortest paths that edge is a part of. Edges with high betweenness centrality are often critical bottlenecks, since their removal can substantially increase path lengths or fragment connections between regions of the network. In what follows, we will use the *average node betweenness centrality* and the *average edge betweenness centrality* which are averaged over all nodes and all edges, respectively.

The longest shortest path length from a node to any other nodes within its component is called its eccentricity. From this, we derive the *LCC average eccentricity*, representing the mean eccentricity across all nodes in the LCC. The minimum eccentricity within the LCC corresponds to its radius. The fraction of nodes in the LCC with an eccentricity equal to the radius is referred to as *FONs in the LCC center*. Conversely, the maximum eccentricity within the LCC determines its diameter. The fraction of nodes in the LCC whose eccentricity equals its diameter is referred to as *FONs in the LCC periphery*. Additionally, the barycenter of the LCC corresponds to the nodes with the lowest total shortest path length to all other nodes. The fraction of nodes thus identified is referred to as *FONs in the LCC barycenter*. Finally, *LCC efficiency* is calculated as the inverse of the shortest path length between each pair of cells in the LCC, providing a measure of the network’s overall efficiency in facilitating connectivity..

Furthermore, we consider measures that capture the tendency of nodes to form tightly connected groups. The *average clustering coefficient* quantifies the prevalence of triadic (three-node) connections, while the *average squares clustering coefficient* extends this concept to tetradic (four-node) structures. *Transitivity* is also measured as the likelihood that a connected triplet of nodes forms a triangle. We identify community structures in the graph using the Girvan-Newman algorithm,^52^ which detects clusters of nodes that are densely connected internally but only loosely connected to other groups. The total *number of communities* provides insight into the network’s modular organization. To further evaluate whether the network exhibits strong intra-community connectivity and sparse inter-community links, we compute *modularity* using the Clauset-Newman-Moore greedy modularity maximization algorithm.^53^ Higher modularity scores indicate that nodes within the same community are more densely interconnected, while connections between different communities remain weaker.

Additional mathematical definitions and methodological details are provided in Supplementary Material, Section C. All computations were performed in Python, primarily using the *NumPy*^54^ and *NetworkX*^55^ libraries.

### 2.10 Classification of maturation stages using graph theoretical features

We analyze the relationships among graph fingerprint measures using Pearson correlation. Each graph fingerprint is represented as an 18-dimensional vector, where each component corresponds to a specific graph-theoretical feature extracted from networks reconstructed from primary rat cortical neuronal cultures at different maturation stages. We compute pairwise correlations between these features across all 192 neuronal network samples, generating a correlation matrix, which is then hierarchically clustered via Clustermap^56^ to group features with similar correlation patterns.

We use a random forest classifier^57^ to identify the most informative features for distinguishing maturation stages of primary rat cortical neuronal cultures. To achieve this, we randomly select 80% of the 192 data vectors and train the classifier on pairs of graph features to predict the maturation stage with the highest precision, as indicated in the confusion matrix. This approach enables us to to accurately classify maturation stages using only a pair of graph measures.

### 2.11 Statistical analysis

For each maturation stage, distributions of graph feature values are extracted, and their means are computed. Welch’s t-test, which does not assume equal variances or equal sample sizes, is then applied to determine whether the mean values differ significantly between two independent maturation stages. Raw p-values were corrected for multiple comparisons using the Benjamini-Hochberg FDR procedure, and significance was assessed using FDR-adjusted q-values. Statistical analyses are performed using the *SciPy* package.^58^ Statistical significance is defined as follows: *p* ≤ 0.05 (*) indicates significance, *p* ≤ 0.005 (**) indicates high significance, and *p* ≤ 0.0005 (***) indicates extreme significance, while results with *p >* 0.05 are considered not significant (n.s.).

## 3 Results

### 3.1 High-performance U-Net segmentation and graph fingerprinting delineate neuronal network structures

The performance of the U-Net models for segmenting cell bodies and neurites from DHM phase images (Section 2.5) is summarized in Fig. 3 A-B. The cell body segmentation model achieved an AUC of 0.98 across the average of 100 ROC curves (Fig. 3. A-1), indicating near-perfect classification, while the neurite segmentation model reached an AUC of 0.91 (Fig. 3 B-1), demonstrating high accuracy. Due to the presence of coherent noise in the background of the phase image and clusters of cells, the F-score, compared to balanced accuracy, reduces the area of predicted objects and decreases false positive rates. Consequently, the optimal thresholds, determined by maximizing the F-score, were 0.3 for cell bodies and 0.59 for neurites. These thresholds are shown in Fig. 3 A-2 and B-2, with representative segmentation results depicted in Fig. 3. A-3 and B-3.

**Fig 3.**
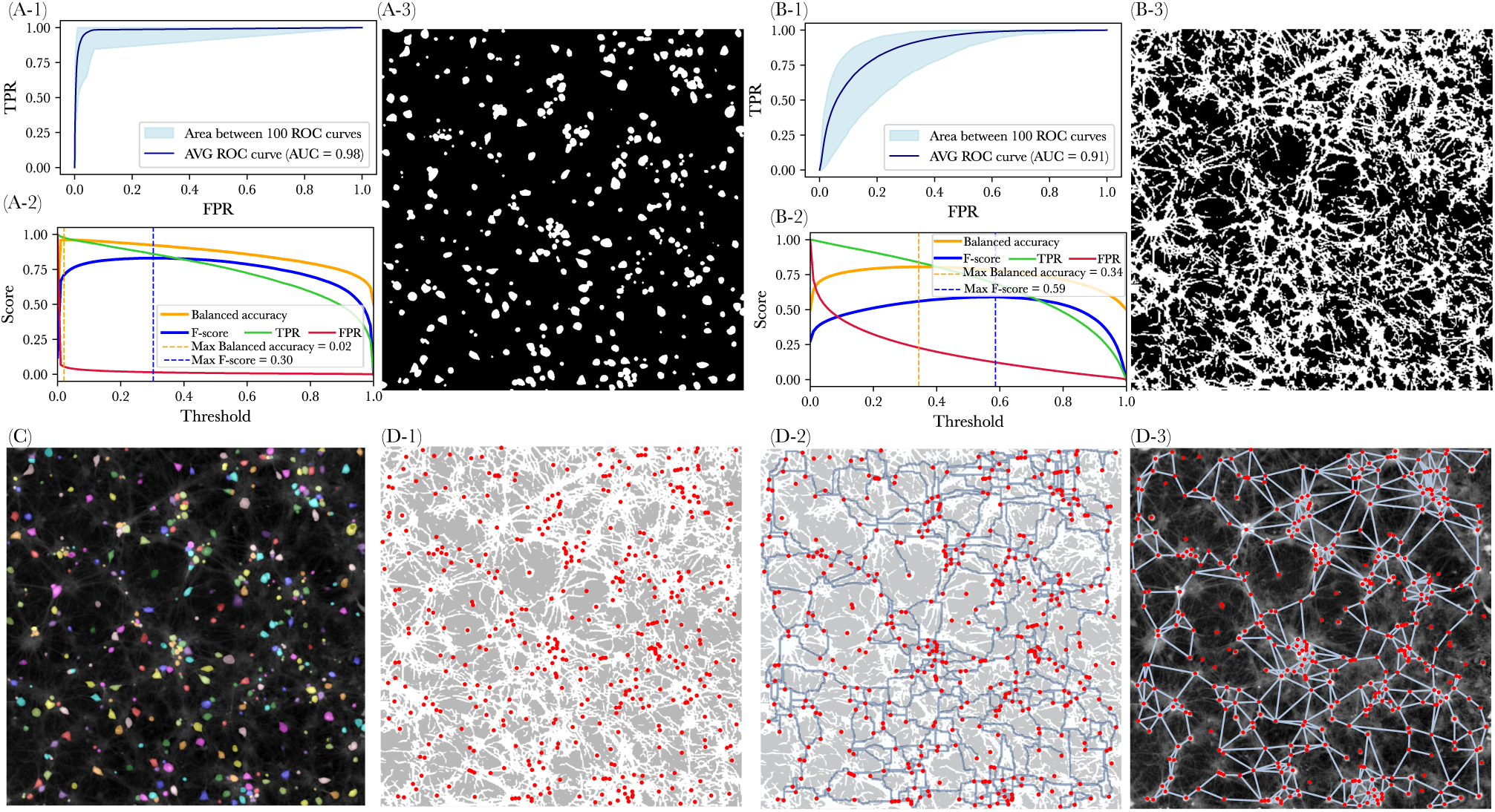
U-Net models performance and infering graph from phase image. **(A)** U-Net segmentation performance for cell bodies: **(A-1)** AUC of the average 100 ROC curves is 98%. **(A-2)** Metrics derived by applying thresholds to the cell body probability map. **(A-3)** Binary mask image with the maximum F-score threshold: 0.3. **(B)** U-Net segmentation performance for neurites: (B-1) AUC of the average 100 ROC curves is 91%. **(B-2)** Metrics derived by applying thresholds to the neurite probability map. **(B-3)** Binary mask image with the maximum F-score threshold: 0.59. **(C)** Separated cell bodies are visualized with the watershed algorithm with distinct colormaps. **(D)** Graph infering of phase image: **(D-1)** Overlaid masks of neurites (white) with red dots marking cell bodies centroids as graph nodes. **(D-2)** Shortest paths (geodesic distances) between cell body pairs shown in light blue. **(D-3)** Graph fingerprint embedded in the phase image, with edges(light blue) representing shortest paths.

In Fig. 3 C, overlapping cell bodies are separated and labeled (Section 2.7). Fig. 3 D-1 displays a combined mask in which cell body centroids, labeled and marked by red dots, are overlaid on the neurite mask. By applying the graph inference procedure to this combined mask, we identify potential shortest paths between pairs of cells through neuronal processes (Fig. 3 D-2, possible paths are shown in light blue) as outlined in Section 2.8. The resulting graph, embedded in the phase image, is illustrated in Fig. 3 D-3, where paths between nodes (cell bodies) are represented by edges. This computational framework is applied to all 192 phase images to extract the detailed structural organization of neuronal networks (Section 2.3).

### 3.2 Graph fingerprint topology undergo significant changes during maturation

Applying our computational framework on the longitudinal dataset revealed distinct changes in graph fingerprint topology throughout the maturation stages of neuronal cell culture. From the full set of 18 graph-derived features, 6 representative features are highlighted in Fig. 4 because they capture the major aspects of network maturation. In each box plot, the diamond denotes the mean, the horizontal pink line the median, the box the standard deviation, the whiskers the minimum and maximum values, and the dots individual measurements. The remaining 12 graph features (see Table 1) reflecting the topological properties of the rat neuronal cell culture graph fingerprints are presented in the Supplementary Material at Section D and in Fig. S1.

**Fig 4.**
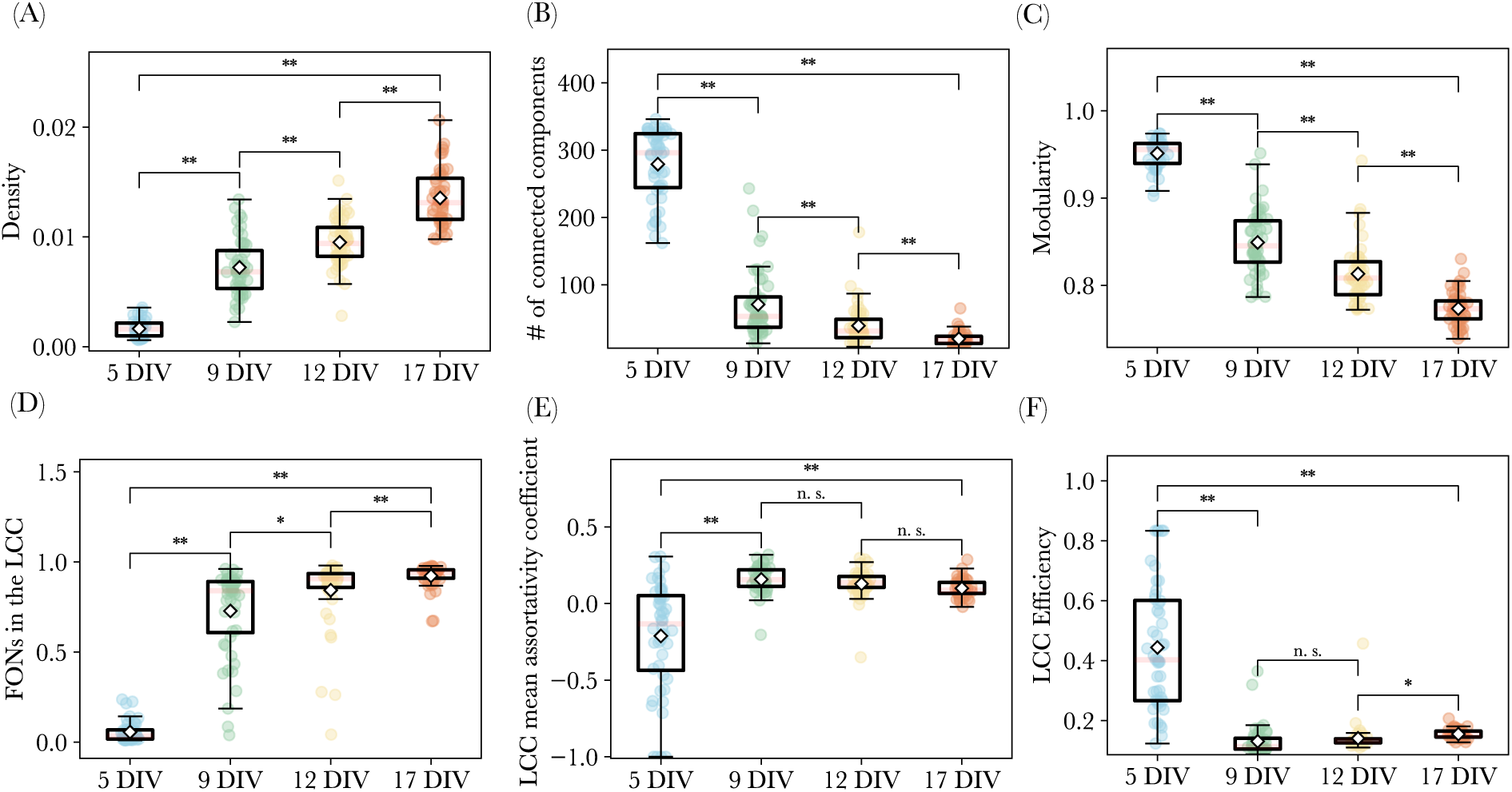
Time evolution of the rat cell culture graph fingerprint topological properties (A-F). **(A)** Graph fingerprint density of rat cell culture. **(B)** # of connected components. **(C)** Modularity score. **D)** FONs of each graph fingerprint contained in its LCC. **(E)** Average assortativity coefficient of LCC **(F)** Global efficiency of the LCC in each graph fingerprint. For all plots, the 192 (48 per DIV) collected FOVs and their corresponding graph fingerprints were used.

From the perspective of graph fingerprint topology, we observe a significant increase in the average density of graph fingerprints (Fig. 4 A; 5 DIV = 0.002, 9 DIV = 0.007, 12 DIV = 0.009, and 17 DIV = 0.013; all statistical comparisons **). This aligns with the decrease in the number of connected components (Fig. 4 B; 5 DIV = 279.1, 9 DIV = 70.9, 12 DIV = 39.1, and 17 DIV = 19.8; all statistical comparisons **), transitioning from numerous isolated cells and groups at earlier stages to a more connected network by 17 DIV. Additionally, the mean modularity score decreases significantly (Fig. 4 C; 5 DIV = 0.95, 9 DIV = 0.85, 12 DIV = 0.81, and 17 DIV = 0.77; all statistical comparisons **), indicating a transition from highly disconnected structures toward networks where connections are formed randomly but are more uniformly distributed spatially.

Furthermore, in Fig. 4 D, the mean FONs in the LCC at 5 DIV is only 6%, increasing to 73% at 9 DIV, 84% at 12 DIV, and 92% at 17 DIV (all statistical comparisons **, and 9 DIV vs. 12 DIV *). The LCC mean assortativity coefficient increases from −0.21 at 5 DIV to 0.16 at 9 DIV, before stabilizing at 0.13 at 12 DIV and 0.10 at 17 DIV, as illustrated in Fig. 4 E (all statistical comparisons **, except 9 DIV vs. 12 DIV and 12 DIV vs. 17 DIV both n.s.). In Fig. 4 F, we note a decline in the LCC efficiency from 0.44 at 5 DIV to 0.13 at 9 DIV, stabilizing at 0.14 at 12 DIV before experiencing a slight increase to 0.15 at 17 DIV (all statistical comparisons **, except 9 DIV vs. 12 DIV n.s., and 12 DIV vs. 17 DIV *).

### 3.3 Maturation of neuronal networks characterized by two key topological properties

To capture a comprehensive view of the various behaviors of graph features over the course of maturation, and to determine how these features co-evolve, we conduct correlation and cluster analyses. These analyses reveal that all the graph features regroup into two highly correlated families, as shown in Supplementary Material Section D, Fig. S1 N. This finding suggests that the maturation of neuronal network structures can be effectively characterized by just two predominant topological properties. To validate this hypothesis and identify the most informative topological properties, we applied a Random Forest classifier to classify maturation stages using graph-derived features (Section 2.10). The classifier achieved an accuracy of 87%, with density and modularity confirmed as the two most discriminative graph features, sufficient to distinguish maturation stages. The accuracy of predicting each stage label is shown as a confusion matrix in Fig. 5 A. The co-evolution of these two features is shown in Fig. 5 B, indicating a relative separation between cultures over maturation. Misclassifications are represented by black circles, from left to right, showing instances in which cell cultures were predicted as 5, 9, 12, 12, and 17 DIV.

**Fig 5.**
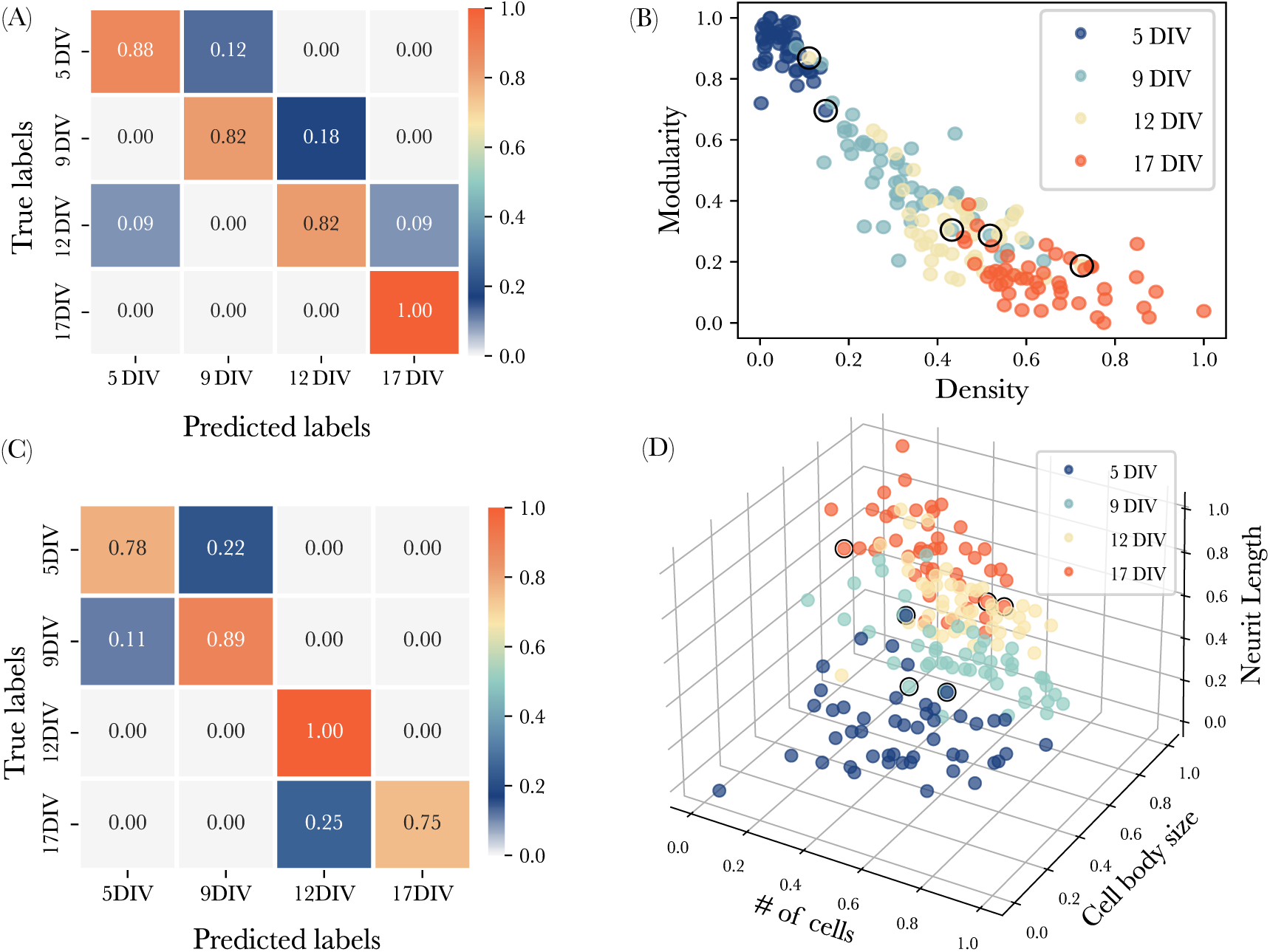
Topological and morphological characterization of neuronal network maturation. **(A)** Confusion matrix showing the performance of a Random Forest classifier in predicting culture maturation stages from two graph-derived features, density and modularity, achieving an accuracy score of 0.87. **(B)** Scatter plot of these two features for all 192 graph fingerprints, with colors indicating maturation stages. Black circles indicate misclassified labels by the Random Forest classifier, which are, from left to right, predicted as 5, 9, 12, 12, and 17 DIV. **(C)** Confusion matrix describing the performance of a multi-layer perceptron (MLP) classifier in classifying morphological properties into culture maturation stages using cell body size, neurite length, and cell number as morphological features, achieving an accuracy score of 0.85. **(D)** Scatter plot of these three features for all 192 phase images, with colors indicating maturation stages. Black circles indicate misclassified labels by the MLP classifier, corresponding to 5 DIV predicted as 9 DIV, 9 DIV predicted as 5 DIV, and 17 DIV predicted as 12 DIV.

In contrast, using pairs of morphological properties corresponding to cell density, neurite length, and cell body area (Supplementary Material Fig. S2) did not exceed 74% accuracy in predicting maturation stages (Supplementary Material Section G and Fig. S3). Considering all three morphological features simultaneously improved classification performance. As shown in the confusion matrix in Fig. 5 C, the resulting classifier achieved an accuracy of 85%. A 3D scatter plot, in which each axis corresponds to one morphological feature, is shown in Fig. 5 D. Misclassifications are represented by circles, showing instances in which 5 DIV is predicted as 9 DIV, 9 DIV as 5 DIV, and 17 DIV as 12 DIV.

## 4 Discussion

In this study, we have taken a significant step forward in exploring the full potential of DHM for analyzing neuronal cell cultures at the network level. Previous research has primarily focused on the structural and functional properties of neurons at the individual level,^1, 59^ often overlooking the complex interactions within the neuronal network. Our work leverages high-performance U-Net segmentation models and a pathfinding algorithm to extract detailed topological and morphological features of neuronal networks from DHM phase images, demonstrating DHM’s ability to provide comprehensive insights into the organization and maturation of neuronal networks.

Our results show that U-Net models achieve high performance in segmenting cell bodies and neurites of DHM phase images, with AUC values of 0.98 and 0.91, respectively. The inferred graph fingerprints provide an effective mathematical model for the structural organization of neuronal networks. Applying our computational framework to a longitudinal dataset of 192 phase images of neuronal networks in culture, evenly collected across four maturation stages (5, 9, 12, and 17 DIV), revealed distinct changes in both topological and morphological properties as the networks mature.

At the topological level, the computational framework was complemented by an array of 18 normalized graph features, inspired by work in Network Neuroscience, to characterize the topology of the graph fingerprints. Our analysis revealed significant changes during neuronal network maturation. For instance, the connected components entropy decreased over time, indicating less compartmentalized and more integrated network structure. In addition, degree assortativity remained close to zero or slightly negative, suggesting little preference for nodes to connect to others of similar degree, consistent with previous studies on cortical networks in vitro.^60, 61^

Moreover, node eccentricity decreased, indicating shorter maximum topological distances, whereas LCC efficiency decreased, reflecting lower average pairwise efficiency within the LCC. These seemingly contrasting trends suggest a reorganization of network topology during maturation, potentially reflecting trade-offs between network integration and wiring organization.^62, 63^

The clustering coefficient increased, suggesting higher network segregation, essential for neuronal development and plasticity.^64–66^ These changes highlight the dynamic nature of neuronal networks, with increased clustering and fewer separated components, possibly reflecting the critical role of GABA in postnatal rat brain development.^67, 68^

At the morphological level, as presented in Supplementary Material Section E, and illustrated in Fig. S2, our analysis showed that cell density remained stable during the initial stages but decreased at later stages. Neurite length and cell body size showed consistent growth, reflecting cellular expansion and increased spacing. This may be linked to the neurotransmitter GABA, which plays a significant role in neuronal development and regulation.^69–71^ For instance, in vitro studies have shown that GABA increases neuronal projection length, neurite branching, and synapse density when applied to embryonic chick cortical and retinal cells.^67, 72^ The maturation GABA functional switch has been reported to occur around the second week in vitro, with most neurons completing the transition by approximately 12 DIV.^73^ This period overlaps with the stage at which several graph-theoretical features changed markedly in the present study. Although this temporal correspondence is intriguing, it remains speculative here because GABAergic maturation was not directly measured. Future studies combining graph-theoretical analysis with direct markers of GABAergic maturation could help clarify this potential relationship.

Building on these topological insights, correlation and clustering analyses of all 18 graph features revealed that two primary features—modularity and density—were sufficient to characterize the maturation process and were the most discriminative descriptors of maturation stage. This was validated using a Random Forest classifier, which achieved an accuracy of 87% in predicting the maturation stages of neuronal cultures. In comparison, three morphological properties—cell density, neurite length, and cell body size— achieved a maximum accuracy of 85% in predicting DIV labels. Despite the greater computational and interpretive complexity of graph-theoretical features, density and modularity alone achieved predictive performance comparable to that obtained using all three morphological descriptors, indicating that neuronal maturation can be effectively captured by a compact topological representation.

## 5 Conclusion

In this study, we developed a computational framework combining DHM, deep-learning-based segmentation, and graph-theoretical analysis to quantitatively characterize neuronal network organization in primary rat cortical cultures. By extracting graph fingerprints from label-free DHM phase images and analyzing a panel of connectomics-inspired descriptors, we demonstrate that the maturation of neuronal networks in vitro is accompanied by reproducible changes in network topology and connectivity.

Beyond conventional cell-level measurements, our approach enables the quantitative characterization of neuronal cultures at the network level. In particular, graph-theoretical descriptors capture the progressive reorganization of network architecture during maturation and provide a compact representation of changes in connectivity and community structure. These findings highlight the potential of combining label-free quantitative imaging with graph analysis to study neuronal networks as integrated systems rather than collections of individual cells.

More broadly, this framework opens new opportunities for pharmacological experiments and drug-screening studies aimed at quantifying the effects of neuroactive compounds on neuronal network organization and maturation.^74^ It may also prove valuable for the study of human induced pluripotent stem cell (hiPSC)-derived neuronal cultures from patients with neuropsychiatric and neurological disorders,^75, 76^ where quantitative and label-free assessment of network phenotypes remains a major challenge. Finally, extending this approach toward functional imaging modalities could provide complementary insights into the relationship between network structure and activity, further advancing our understanding of neuronal network physiology and pathology.^77, 78^

## 6 Code and Data Availability

The source code and trained models for the two U-Net pipelines used for cell body and neurite segmentation are publicly available through Zenodo.^79^ The data supporting the findings of this study are available from the corresponding author upon reasonable request.

## 7 Acknowledgements

The authors sincerely thank Céline Larivière-Loiselle, Marie-E^’^ ve Crochetière, and Johan Chaniot for their invaluable assistance and support during this research. This work was supported by: the Fonds d’accélération des collaborations en santé du Québec (Alliance Neuro-CERVO), a pro-gram coordinated by the Consortium québécois sur la découverte du médicament (ZY, EB, AA, PM, PD); the Natural Sciences and Engineering Research Council of Canada (AA, PM, PD); and the Sentinelle Nord program of Université Laval, funded through the Canada First Research Excellence Fund (AA, PM, PD). We acknowledge Calcul Québec and Digital Research Alliance of Canada for their technical support and computing infrastructures.

## 8 Author contributions statement

ZY, EB, JL, AA, PM, and PD designed the project. ZY conducted all experiments, reconstructed DHM phase images with a digital holography library from MH, and coded most of the computational framework with help from MM, AA, and PD. All authors analyzed the results. EB, and PM supervised the experimental part of the project while AA and PD supervised theory. ZY, EB, and PD wrote the first draft, and all authors reviewed the manuscript.

## Supplementary Material

### A Labeling phase images with existing methods

Before developing our segmentation approach, we evaluated existing methods for cell body and neurite identification. In the case of cell bodies, we tested Cellpose’s cyto model^23^ with its graphical user interface. Although this model is widely used for cell body segmentation, we found that it did not perform well on our DHM phase images due to several shortcomings. First, because DHM phase images were not included in the Cellpose training dataset, the model fails to account for coherent noise. In addition, since the QPS is both non-specific and axially cumulative, it is difficult to distinguish neurite branches near cell bodies in DHM phase images.^80^ During this project, the second version of Cellpose was also released.^81^ Although Cellpose 2.0 allows training on specific image types and tasks, its architecture closely resembles the model we were already developing. Thus, we determined that adopting this model would not provide significant advantages over our own and opted not to use it.

To segment neuronal processes, we tested both semi-automatic and automatic methods. We used the Trainable Weka Segmentation plugins in Fiji, a machine learning tool capable of training a classifier with a limited number of manual annotations and then performing automatic segmentation on the remaining data. In addition, we tried to segment neuronal processes using DeepNeurite. However, these approaches exhibited limitations in handling DHM phase images. Weka struggled to capture all regions of interest and remove background noise, likely due to the high feature complexity, coherent noise, and variations in connection width within the phase images. Similarly, DeepNeurite struggled to delineate neuron branches of varying width. This is primarily because its training dataset consisted primarily of fluorescence imaging data of neuronal processes with relatively consistent widths. In addition, the training data does not account for the unique structural characteristics of cell culture growth, which can change significantly as the culture matures. Therefore, we did not adopt either approach for our application. These limitations underscore the need to tailor segmentation methods to the unique characteristics of DHM phase images.

### B U-Net architecture

The U-Net model was implemented in Python and adapted for the Keras and TensorFlow libraries. More specifically, it allows for padded convolutions, adjustment of network depth and width, drop-out rate, and batch normalization. The U-net structure consists of both down-sampling and up-sampling paths. The down-sampling path consists of four blocks, each containing two 3×3 convolutional layers, followed by a ReLU activation function and a batch normalization layer. A dropout layer is inserted between the convolutional layers for generalization, while zero padding ensures consistent channel dimensions across layers. Downsampling is achieved with 2×2 max-pooling layers, resulting in a twofold reduction in image size. A bottleneck convolutional block connects the down-sampling and up-sampling paths. The upsampling path contains four blocks, each with two 3×3 convolutional layers, followed by a ReLU activation function and batch normalization. Spatial upsampling is performed using transposed convolutional layers, doubling the image resolution. Skip connections via concatenation integrate low-level and high-level features to improve segmentation accuracy. The final convolutional layer reduces the 32 feature channels to a single channel using a sigmoid activation function, producing a grayscale output image. The U-Net models for cell body and neurite segmentation were trained on an M1 MacBook Air with 8 GB of memory.

### C Graph features

This section provides definitions for all graph features used in our study, all of which have been previously applied in network neuroscience, with most adapted from Ref.^30^ The graphs generated in this study are unweighted, undirected, and contain no self-loops, corresponding to the notion of a simple graph in graph theory.^82^

### Preliminary definitions and notation

A simple graph is an ordered pair *G* = (*V, E*), where *V* is a finite set of nodes, and *E* is a set of edges, each represented as an unordered pair of distinct nodes {*i, j*} with *i, j* ∈ *V* and *i* ≠*j*. Thus, *E* is a subset of the set of all unordered pairs of distinct elements of *V*. The number of nodes is *N* = |*V* |, and the number of edges is *M* = |*E*|, where |*S*| represents the cardinality of a set *S*. The **adjacency matrix** of *G* is the *N* × *N* matrix *A* = (*a_ij_*)*_i,j_*_∈_*_V_* such that *a_ij_* = 1 if {*i, j*} ∈ *E* and *a_ij_* = 0 otherwise. For a simple graph, the adjacency matrix is symmetric and has zeros on the diagonal. The **degree** of a node *i*, denoted *k_i_*, is the number of nodes to which it is directly connected, i.e., *k_i_* = [inlnine]. A **path** of length *ℓ* between nodes *i* and *j* is a sequence of distinct nodes (*i* = *v*_0_*, v*_1_*, …, v_ℓ_* = *j*) such that {*v_m_, v_m_*_+1_} ∈ *E* for all *m* ∈ {0*, …, ℓ* − 1}. The **shortest path length** *d*(*i, j*) is the minimal number of edges in any path between *i* and *j*, with *d*(*i, j*) = ∞ if no such path exists. A **connected component** is a maximal subset of nodes in which every pair is connected by a path. The **largest connected component (LCC)** is the component with the most nodes, denoted by *N*_LCC_. The **topology** of a graph refers to the arrangement of its nodes and edges, describing its overall structure. In a simple graph, topology is fully determined by the adjacency matrix. Graph features are topological invariants, meaning they quantify structural properties that remain unchanged under node relabeling. If two simple graphs do not share the same features, then they are necessarily topologically distinct. The number of vertices *N*, the number of edges *M*, the set of degrees {*k_i_* | *i* ∈ *V* }, and the set of shortest path lengths {*d*(*i, j*) | *i, j* ∈ *V* } are basic topological invariants. More refined graph features are defined below.

### 1. Number of connected components

The set of vertices *V* admits a unique disjoint partition into *k* connected components, meaning *V* = *V*_1_ ∪ · · · ∪ *V_k_* and *V_c_* ∩ *V_d_*= ∅, for all 1 ≤ *c < d* ≤ *k*, where each *V_c_* is a connected component. The number of connected components is denoted by *k*. If *N_c_* denotes the size of component *V_c_*, then the total number of nodes Satisfies 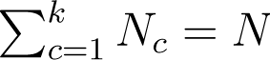.

### 2. Fraction of nodes (FONs) in the largest connected component (LCC)

This is the total number of nodes in the LCC, *N*_LCC_, divided by total number of nodes in the graph:

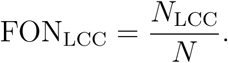

### 3. Connected components entropy

This is the normalized Shannon entropy related to the distribution of nodes across the connected components of the graph:

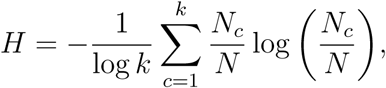

where log denotes the logarithm in base 2. The normalization factor ensures that 0 *< H* ≤ 1. The entropy reaches its maximum value when all connected components contain the same number of nodes, meaning 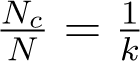 for all *c* and for any *k*.

### 4. Density

The edge density of the graph, defined as the fraction of observed edges relative to the total number of possible edges in a fully connected graph:

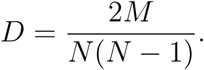

### 5. LCC mean associativity coefficient

The assortativity coefficient of a simple graph *G* = (*V, E*) is the Pearson correlation coefficient of the vertex degrees, *k_i_* for *i* ∈ *V*, between pairs of connected vertices:

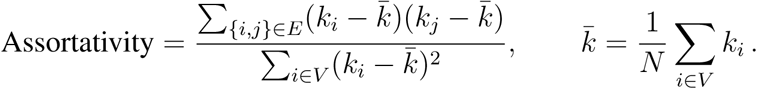

A positive assortativity coefficient indicates that nodes tend to connect to others with similar degrees. When the calculation is restricted to edges within the largest connected component (LCC), we obtain the LCC mean assortativity coefficient.

### 6. LCC average distance

The average distance in a graph is the mean shortest path length between all pairs of nodes:

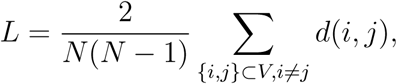

where *d*(*i, j*) is the shortest path length between nodes *i* and *j*, as previously defined. If the graph is disconnected, *d*(*i, j*) can be infinite, making *L* ill-defined. To ensure a finite value, the computation is restricted to the LCC, where all nodes are mutually reachable. The resulting quantity is referred to as the LCC average distance.

### 7. Node betweenness centrality

The betweenness centrality of a node quantifies its importance in shortest path routing within the network. It is defined as the fraction of all shortest paths that pass through a given node. The betweenness centrality of node *i* is given by:

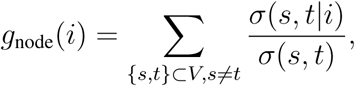

where *σ*(*s, t*) is the total number of shortest paths between nodes *s* and *t*, and *σ*(*s, t*|*i*) is the number of those paths that pass through *i*. If *i* does not lie on any shortest path between *s* and *t*, then *σ*(*s, t*|*i*) = 0, ensuring that *g*_node_(*i*) remains well-defined. The average node betweenness centrality is given by:

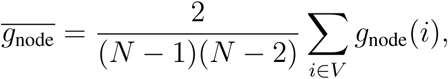

where (*N* − 1)(*N* − 2)*/*2 is the number of distinct reachable node pairs in an undirected graph.

### 8. Edge betweenness centrality

The betweenness centrality of an edge *e* quantifies its role in shortest path routing within the network. It is defined as:

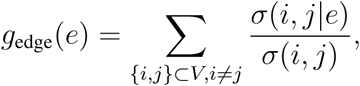

where *σ*(*i, j*) is the number of shortest paths between nodes *i* and *j*, and *σ*(*i, j*|*e*) is the number that pass through *e*. If *e* is not part of any shortest path between *i* and *j*, then *σ*(*i, j*|*e*) = 0. The average edge betweenness centrality is:

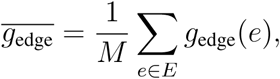

where *M* is the total number of edges in the graph.

### 9. LCC average eccentricity

The eccentricity of node *i* is the maximal shortest path length between *i* and any other node:

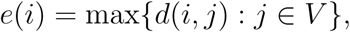

where *d*(*i, j*) is the shortest path length between nodes *i* and *j* The average node eccentricity of a graph is

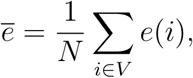

The **radius** *r* of a graph is the minimum eccentricity, min{*e*(*i*)|*i* ∈ *V* }, while the **diameter** *d* is the maximum eccentricity, max{*e*(*i*)|*i* ∈ *V* }. When a graph is disconnected, meaning that it has more than one connected component, some eccentricities are infinite. To avoid this, we restrict the calculations to the LCC, defining the LCC average eccentricity as:

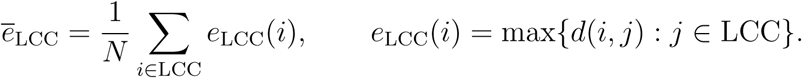

### 10. FONs in the LCC center

The center of a graph is the set of nodes whose eccentricity is equal to the graph radius: Center(*G*) = {*i* ∈ *V* | *e*(*i*) = *r*}. When restricted to the largest connected component (LCC), the FONs in the LCC center is the fraction of nodes in the LCC that belong to its center:

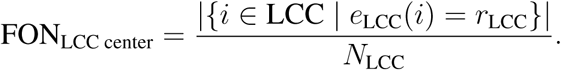

### 11. FONs in the LCC periphery

The periphery of a graph is the set of nodes whose eccentricity is equal to the graph’s diameter: Periphery(*G*) = {*i* ∈ *V* | *e*(*i*) = *d*}. When restricted to the largest connected component (LCC), the FONs in the LCC periphery is the fraction of nodes in the LCC that belong to its periphery:

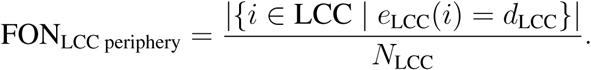

### 12. FONs in the LCC barycenter

The barycenter of a connected graph is the subgraph induced by the set of nodes that minimize the total shortest path distance to all other nodes:

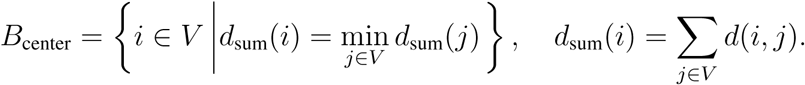

When restricted to the largest connected component (LCC), the FONs in the LCC barycenter is the fraction of nodes in the LCC that belong to its barycenter:

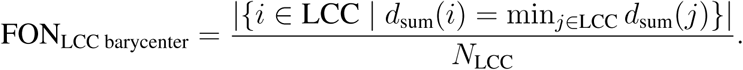

### 13. LCC efficiency

The global efficiency of a graph measures how efficiently information might be exchanged over its network structure. It is defined as:

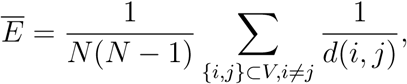

where *d*(*i, j*) is the shortest path length between nodes *i* and *j*.

When the graph is disconnected, some shortest path lengths *d*(*i, j*) are infinite, making efficiency ill-defined. To ensure a finite value, the calculation is restricted to the LCC, yielding the LCC efficiency:

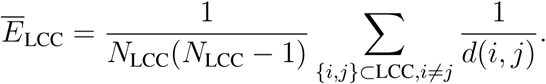

### 14. Average clustering coefficient

The clustering coefficient of a node *i*, denoted *C_i_*, quantifies the likelihood that its neighbors are also connected to each other, forming a triangle:

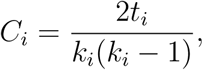

where *t_i_* is the number of triangles that include node *i*, and *k_i_* is its degree. If *k_i_ <* 2, then *C_i_* is set to 0. The average clustering coefficient of the graph is given by:

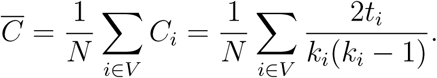

This measure quantifies the tendency of a node’s neighbors to be connected, indicating the presence of tightly-knit clusters in the network.

### 15. Average squares clustering coefficient

This coefficient quantifies the tendency of a node’s neighbors to form four-node cycles (squares). It is defined as the average ratio of the number of observed squares involving a node to the total number of possible squares that could include that node:

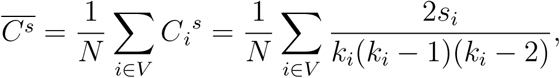

where *C_i_^s^* is the square clustering coefficient of node *i*, and it is set to 0 if *k_i_ <* 3. The quantity *s_i_* represents the number of squares passing through node *i*, given by:

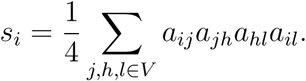

### 16. Transitivity

This measure quantifies the overall tendency of a graph to form triangles, capturing the global density of closed triplets. It is defined as the fraction of all possible triplets (connected node pairs with a common neighbor) that actually form triangles: number of closed triplets

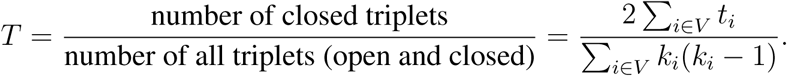

Transitivity is closely related to the average clustering coefficient. However, unlike the latter coefficient, transitivity does not assign a coefficient to individual nodes but instead provides a single global measure for the entire network.

### 17. Number of communities

A **community** (also called a module or cluster) in a graph is a subset of nodes that are more densely connected internally than to nodes outside the subset. In this study, we detect communities using the Girvan-Newman algorithm,^83^ which iteratively removes edges with the highest betweenness centrality to reveal hierarchical community structures. The number of communities, denoted by *q*, is the total number of detected clusters in the network. A higher *q* indicates a more fragmented or modular structure, while a lower *q* suggests stronger global connectivity.

### 18. Modularity

A **community structure** of a graph is a partition of *V* into *q* nonoverlapping subsets (communities or modules), such that edges are more densely distributed within a community than between communities. The **modularity** *Q* quantifies the quality of this partition by comparing the fraction of edges within modules to the expected fraction in a randomized network. It is given by

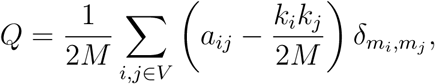

where *M* is the total number of edges, *a_ij_*is the adjacency matrix entry, *k_i_*is the degree of node *i*, and *m_i_* ∈ {1*, …, q*} denotes the module to which node *i* belongs. The term *δ_m__i,mj_* is equal to 1 if nodes *i* and *j* belong to the same module and 0 otherwise. Higher values of *Q* usually indicate stronger modular organization, with a clear separation between communities. In this study, modularity is computed using the Clauset-Newman-Moore greedy modularity maximization algorithm,^53^ which iteratively merges communities in a bottom-up fashion to maximize *Q*. Starting with each node as its own community, the algorithm greedily merges the pair that yields the largest modularity increase until no further improvement is possible, returning the final partition and its modularity value.

The list of graph features presented here is far from exhaustive. Numerous additional metrics are widely used in Network Neuroscience,^30, 33–35^ and new ones continue to emerge,^84–86^ including measures designed to detect low-dimensional (low-rank) structure in graphs.^87^ While we tested many such metrics, we retained only those that exhibited significant variance across the entire dataset of graph fingerprints, ensuring their relevance for characterizing maturation changes.

### D Normalization and aggregation of graph features

To obtain a global view of how graph features vary across the entire dataset of graph fingerprints, we normalized each feature across all samples, rescaling them so that the minimum value is 0 and the maximum is 1. We then aggregated the normalized features into a unified representation, as shown in Fig. S1-A. On the horizontal axis, we assigned the names of the graph features, while the vertical axis represents the ordered graph IDs starting from 5 DIV to 17 DIV. Each horizontal vector represents a single graph fingerprint with all features. Each feature that is the vertical vector of Fig. S1-A is shown in Fig. S1-B-M. Figure S1-M shows the non-normalized features to provide insight into the values of the number of communities over time. The mean values of the horizontal vectors, along with their interpretations, are summarized below:

- Connected components entropy (Fig. S1. B): The entropy decreases significantly from 5 DIV = 0.93 to 9 DIV = 0.35, 12 DIV = 0.23, and 17 DIV = 0.16; (all statistical comparisons**). This indicates a reduction in the variability of disconnected components as the neuronal network matures, suggesting increasing connectivity.
- LCC average distance (Fig. S1. C): The average distance drops sharply from 5 DIV = 0.28 to 9 DIV = 0.06, 12 DIV = 0.04, and 17 DIV = 0.04; (all statistical comparisons **, except 9 DIV vs. 12 DIV and 12 DIV vs. 17 DIV both ns). This reflects that neurons within the LCC become more closely interconnected over time, consistent with maturation.
- Average node betweenness centrality (Fig. S1. D): Values increase from 5 DIV = 0.000 to 9 DIV = 0.018, 12 DIV = 0.021, and 17 DIV = 0.025; (all statistical comparisons **, and 12 DIV vs. 17 DIV *), indicating that neurons increasingly occupy central positions along shortest paths within the network as it matures.
- Average edge betweenness centrality (Fig. S1. E): A gradual increase is observed from 5 DIV = 0.000 to 9 DIV = 0.013, 12 DIV = 0.014, and 17 DIV = 0.015; (all statistical comparisons **, except 9 DIV vs. 12 DIV and 12 DIV vs. 17 DIV both ns), suggesting that certain connections increasingly serve as topological bridges between different parts of the network.
- LCC Average Eccentricity (Fig. S1. F): Eccentricity decreases from 5 DIV = 0.47 to 9 DIV = 0.11, 12 DIV = 0.09, and 17 DIV = 0.07; (all statistical comparisons **, except 9 DIV vs. 12 DIV and 12 DIV vs. 17 DIV both ns). This shows that the maximum distance from each node to the farthest node in the LCC diminishes as the network becomes denser.
- FONs in the LCC center (Fig. S1. G): The FONs in the LCC center decreases from 5 DIV = 0.151 to 9 DIV = 0.015, 12 DIV = 0.016, and 17 DIV = 0.013; (all statistical comparisons **, except 9 DIV vs. 12 DIV and 12 DIV vs. 17 DIV both ns), suggesting a random structure.
- FONs in the LCC Periphery (Fig. S1. H): FONs in the periphery drop from 5 DIV = 0.280 to 9 DIV = 0.018, 12 DIV = 0.019, and 17 DIV = 0.018; (all statistical comparisons **, except 9 DIV vs. 12 DIV and 12 DIV vs. 17 DIV both ns), reflecting the reduced influence of peripheral nodes as the network consolidates.
- FONs in the LCC Barycenter (Fig. S1. I): Values decline from 5 DIV = 0.111 to 9 DIV = 0.006, stabilizing at 12 DIV = 0.006, and 17 DIV = 0.004; (all statistical comparisons **, except 9 DIV vs. 12 DIV and 12 DIV vs. 17 DIV both ns). This suggests that fewer nodes are in the geometric center of the LCC, consistent with increased network centralization.
- Average Clustering Coefficient (Fig. S1. J): Clustering increases from 5 DIV = 0.05 to 9 DIV = 0.24, 12 DIV = 0.28, and 17 DIV = 0.32; (all statistical comparisons **), indicating that neighboring nodes form increasingly tight clusters as the network develops.
- Average Square Clustering (Fig. S1. K): Square clustering rises from 5 DIV = 0.01 to 9 DIV = 0.05, 12 DIV = 0.07, and 17 DIV = 0.08; (all statistical comparisons **), reflecting the formation of higher-order structural motifs in the network.
- Transitivity (Fig. S1. L): Transitivity increases from 5 DIV = 0.23 to 9 DIV = 0.31, remaining stable at 12 DIV = 0.31, and rising slightly at 17 DIV = 0.33; (all statistical comparisons **, except 9 DIV vs. 12 DIV ns, and 12 DIV vs. 17 DIV *). This reflects the growing tendency of the network to form tightly-knit groups.
- Number of Communities (Fig. S1. M): The number of communities increases from 5 DIV = 158 to 9 DIV = 337, decreases slightly at 12 DIV =320, and further reduces at 17 DIV = 273; (all statistical comparisons **, except 9 DIV vs. 12 DIV ns), suggesting that neuronal network maturation is accompanied by the formation of an increasingly structured community organization during early stages, followed by partial consolidation at later stages.

It is important to note that connected components and communities characterize distinct aspects of network organization. Connected components correspond to disconnected graph fragments, whereas communities correspond to groups of nodes that are more densely connected to each other than to the rest of the graph. Thus, the decrease in the number of connected components indicates that initially separated graph fragments progressively merge into larger connected structures. In contrast, the transient increase in the number of detected communities suggests that the internal organization of these larger structures becomes increasingly differentiated during early maturation, before partially consolidating at later stages.

**Fig S1.**
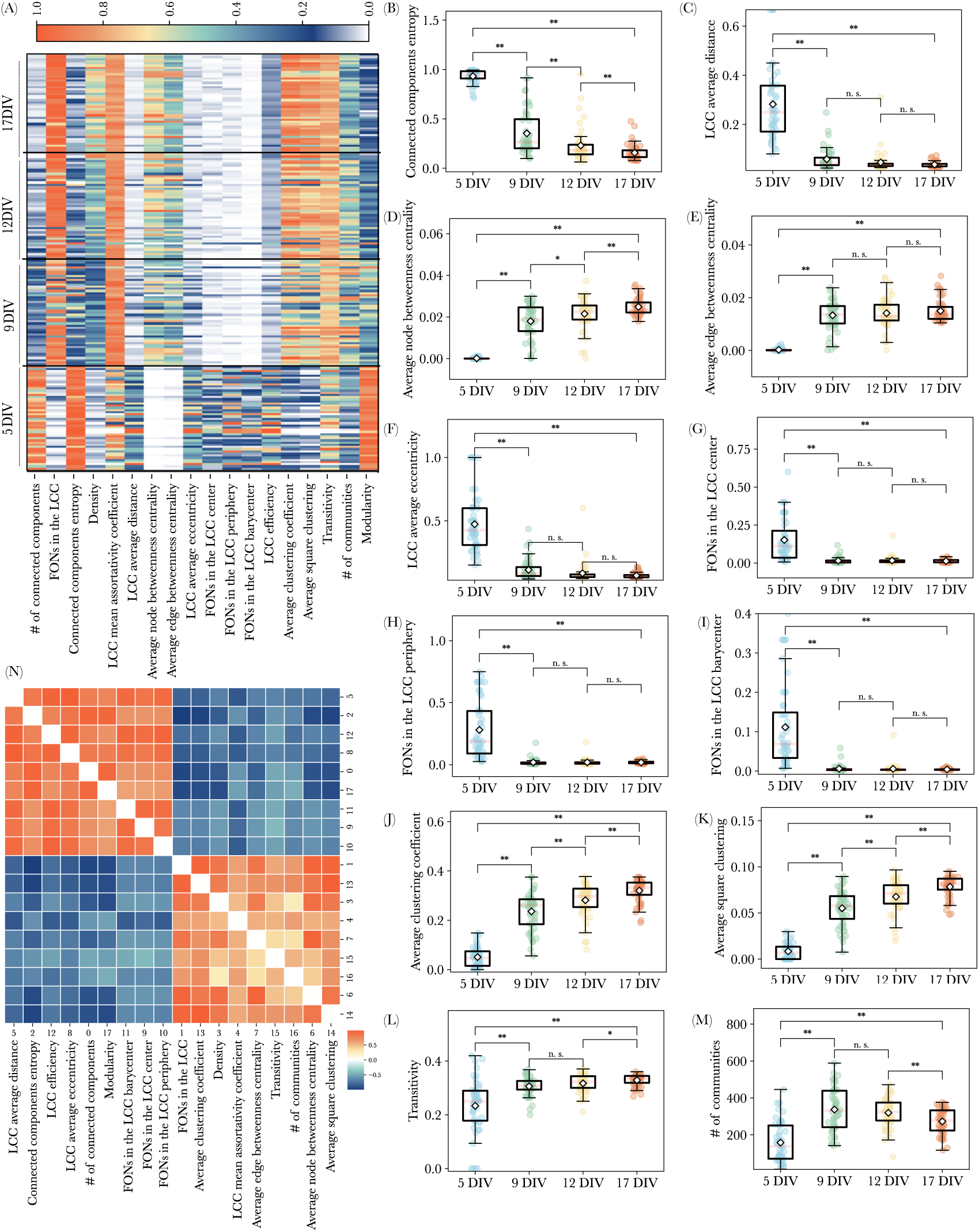
Time evolution of the rat cell culture graph fingerprint topological properties (A-M) and correlation matrix (N). **(A)** 18 normalized graph features of 192 graph fingerprints. Changes from 5 to 9, 12, and 17 DIVs for: **(B)** Connected components entropy, **(C)** LCC average distance, **(D)** Average node betweenness centrality, **(E)** Average edge betweenness centrality, **(F)** LCC average eccentricity, **(G)** FONs in the LCC center, **(H)** FONs in the LCC periphery, **(I)** FONs in the LCC barycenter, **(J)** Average clustering coefficient, **(K)** Average square clustering, **(L)** Transitivity, **(M)** Number of communities detected by the Girvan–Newman algorithm. **(N)** Cluster map of the normalized measures correlation matrix.

### E Neuronal morphology undergo significant changes during maturation

To complement the analysis of graph features, we examined changes in neuronal morphology across maturation stages. Three key morphological properties–cell density, cell body area, and neurite length–were extracted from binary mask images obtained via U-Net segmentation, using the optimal threshold values determined in Section 3.1.

In terms of neuronal morphology, the number of cells within an area of 1*mm*^2^, i.e., cell density, remains relatively stable between 5 DIV (501.9) and 9 DIV (468.4) but shows a significant decrease thereafter, dropping to 412.2 at 12 DIV and further to 335.6 at 17 DIV (Fig.S2-A, all statistical comparison *, except 5 DIV vs. 9 DIV ns). Concurrently, neurite length increases from 25.5 *µm*

### F Optimization and performance validation of Random Forest for maturation stage prediction using graph features

To determine the most effective classification method for predicting maturation stages based on graph fingerprints, we tested multiple machine learning models,, including Logistic Regression, Support Vector Classification (SVC), k-nearest neighbors (k-NN), Decision Tree, Multi-layer Perceptron (MLP), and Random Forest, with default settings to predict the stage of maturation from 192 graph fingerprints features.

To ensure robustness and generalizability, we employed k-fold cross-validation, a standard resampling technique that splits the dataset into k subsets (folds), iteratively training on k-1 folds while testing on the remaining fold. The process is repeated k times, and the average performance metrics are reported.

Among the classifiers, Random Forest exhibited superior generalization ability and robustness in classifying the array of features for DIV compared to the other models. The cross-validation results of these classifiers are summarized in Table S1.

**Table S1.** Cross-validation (CV) results for different classifiers in predicting maturation stages based on graph fingerprints. Accuracy, recall, and macro-averaged F1-score are reported as evaluation metrics.

| Model | CV Accuracy | CV Recall | CV Macro F1-Score |
| --- | --- | --- | --- |
| Logistic Regression | 0.75 | 0.78 | 0.75 |
| SVM | 0.75 | 0.76 | 0.74 |
| k-NN | 0.77 | 0.79 | 0.77 |
| Decision Tree | 0.71 | 0.74 | 0.70 |
| MLP | 0.79 | 0.79 | 0.79 |
| Random Forest | 0.82 | 0.82 | 0.79 |

### G Comparative evaluation of classifiers for maturation stage prediction using morphological features

In Section 3.3, we demonstrated that a small subset of graph features–specifically, density and modularity–was sufficient to predict the maturation stages of primary rat cortical neuronal cultures. Here, we assess whether morphological features alone can achieve comparable predictive performance.

To this end, we trained and evaluated multiple classifiers using three key morphological properties: neurite length, number of cells, and cell body size. As in the previous section, we conducted cross-validation cross-validation (CV) with various models, including Logistic Regression, Support Vector Classification (SVC), k-nearest neighbors (k-NN), Decision Tree, Multi-layer Perceptron (MLP), and Random Forest. Among them, the MLP classifier achieved the highest scores across all evaluation metrics, as shown in Table S2.

**Table S2.** Cross-validation (CV) results for different classifiers in predicting maturation stages based on morphological features. Accuracy, recall, and macro-averaged F1-score are reported as evaluation metrics.

| Model | CV Accuracy | CV Recall | CV Macro F1-Score |
| --- | --- | --- | --- |
| Logistic Regression | 0.71 | 0.73 | 0.70 |
| SVM | 0.74 | 0.75 | 0.74 |
| k-NN | 0.70 | 0.71 | 0.71 |
| Decision Tree | 0.67 | 0.68 | 0.65 |
| MLP | 0.77 | 0.78 | 0.77 |
| Random Forest | 0.75 | 0.74 | 0.72 |

To predict maturation stages, we test the MLP model with pairs of morphological features and calculate the confusion matrix for the pairs listed in Fig. S3. A, B, C, 1 and 2. Using the pair of neurite length and the number of cells, we achieve an accuracy of 0.71 for the MLP. For the pairs of the number of cells and cell body size, and neurite length and cell body size, we achieve an accuracy of 0.74 for the MLP.

**Fig S2.**
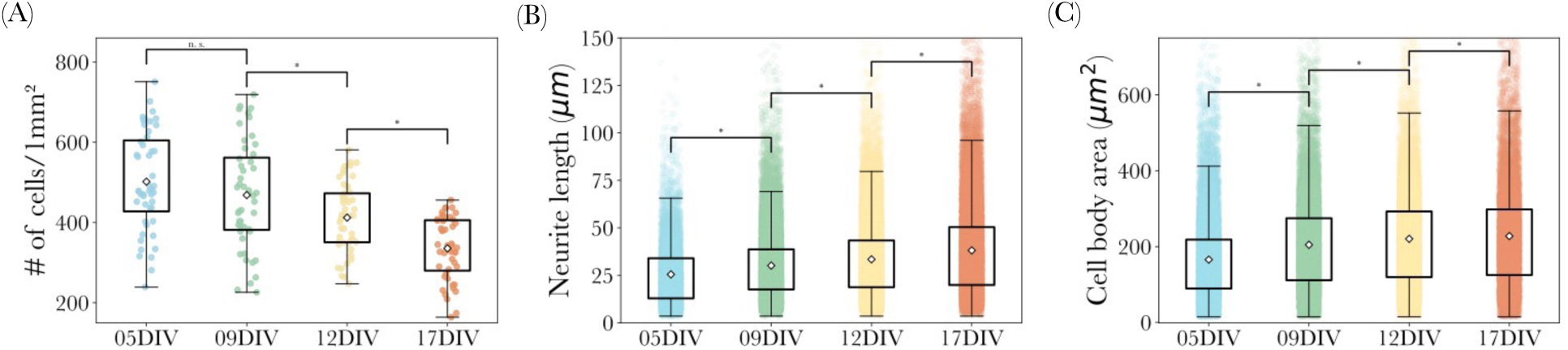
Time evolution of the rat cell culture morphological properties, from 5 to 9, 12, and 17 DIV. **(A)** Density of cells within the FOV of the phase image, measured within an area of 1*mm*^2^. **(B)** Neurite length in *µm* in each FOV. **(C)** Cell body area in *µm*^2^. at 5 DIV to 30.1 *µm* at 9 DIV, 33.4 *µm* at 12 DIV, and 38.1 *µm* at 17 DIV, reflecting neurite outgrowth (Fig.S2-B, all statistical comparison *). The average cell body area similarly expands from 165.6 *µm*^2^ at 5 DIV to 204.7 *µm*^2^ at 9 DIV, 220.4 *µm*^2^ at 12 DIV, and 227.7*µm*^2^ by 17 DIV, illustrating cellular growth (Fig. S2-C, all statistical comparison *).

**Fig S3.**
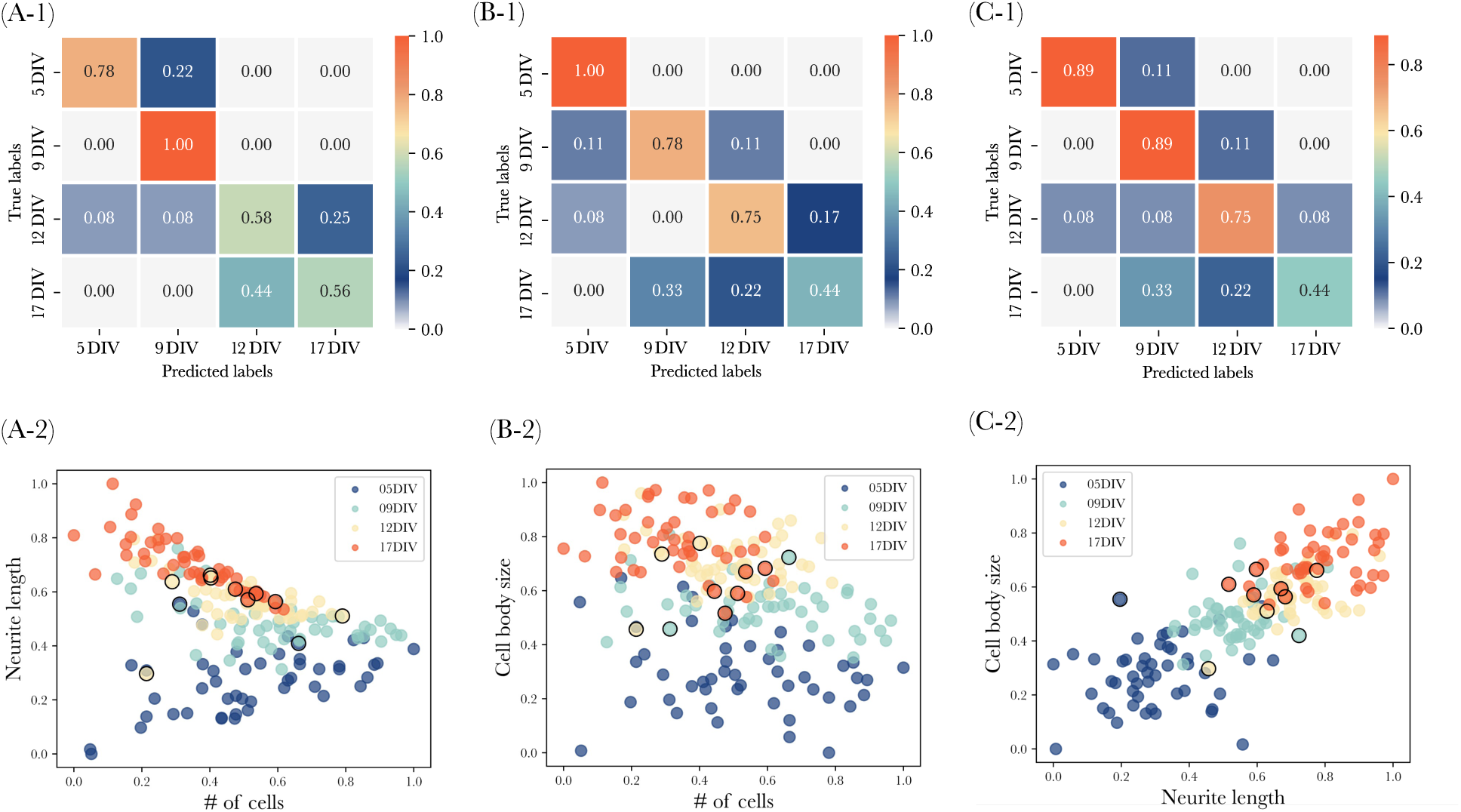
Morphological characterization of neuronal network maturation. **(A-1)** Confusion matrix for MLP classifier performance using neurite length and # of cells, achieving 0.71 accuracy. **(A-2)** Scatter plot of neurite length and # of cells, with colors indicating maturation stages and black circles marking misclassifications, where 5 DIV is predicted as 9 DIV, 12 DIV is predicted as 5 and 17 DIV, and 17 DIV is predicted as 12 DIV. **(B-1)** Confusion matrix for MLP classifier performance using cell body size and # of cells, with 0.74 accuracy. **(B-2)** Scatter plot of cell body size and # of cells, with colors indicating maturation stages and black circles marking misclassifications, where 9 DIV is predicted as 5 and 12 DIV, 12 DIV is predicted as 5 and 17 DIV, and 17 DIV is predicted as 9 and 12 DIV. **(C-1)** Confusion matrix for MLP classifier performance using cell body size and neurite length, with 0.74 accuracy. **(C-2)** Scatter plot of ell body size and neurite length, with colors indicating maturation stages and black circles marking misclassifications, where 5 DIV is predicted as 9 DIV, 9 DIV is predicted as 12 DIV, 12 DIV is predicted as 5, 9 and 17 DIV, and 17 DIV is predicted as 9 and 12 DIV.

